# Human macrophages secrete both interferon α and interferon β protein during infection with Mycobacterium tuberculosis

**DOI:** 10.1101/2025.07.17.665294

**Authors:** Gina Leisching, Donal Cox, Vincent Bondet, Anjali S Yennemadi, Seán Donohue, Darragh Duffy, Joseph Keane

## Abstract

Mycobacterium tuberculosis (Mtb) infection activates type I interferons (IFNs) which are crucial mediators of tuberculosis (TB) pathogenesis. Despite assumptions that IFNα and IFNβ are secreted by macrophages, direct protein quantification in primary human monocyte-derived macrophages is surprisingly lacking. Here, we demonstrate measurable IFNα and IFNβ secretion by MDMs infected with both virulent (H37Rv) and attenuated (H37Ra) Mtb strains as early as 48 h post-infection, with levels persisting at 120 h. These findings challenge existing assumptions about type I IFN kinetics and highlight the importance of timing in experimental designs and provides a foundation for exploring their role in host-pathogen interactions.

## Background

*Mycobacterium tuberculosis* (Mtb) is a pathogen that preferentially infects macrophages and establishes a permissive niche for replication, driving pathogenesis in the host. Upon infection, macrophages adopt a proinflammatory phenotype characterized by the secretion of various cytokines, which the literature suggests may include type I interferons (IFNs). While transcriptional evidence indicates that macrophages engage type I IFN signaling during Mtb infection[1], it remains unclear whether this translates into the actual release of IFNα and IFNβ proteins in human macrophages.

IFNα and IFNβ have attracted significant attention due to their well-established link to tuberculosis (TB) pathogenesis. Progression to active TB has been correlated with heightened type I IFN activity, and elevated interferon stimulated genes have been widely reported in the blood of TB patients [2]. Intriguingly we failed to detect elevated plasma IFNα or IFNβ levels using ultrasensitive digital ELISA in patients with active TB as compared to latently infected controls, despite the presence of a strong ISG signature [3]. This raised the potential hypothesis that IFNα/β is produced locally in the infected tissue where it activates cells in circulation but is not secreted systemically [3]. Despite these different lines of evidence, a fundamental question regarding the macrophage host response to Mtb infection remains: do primary human monocyte-derived macrophages (MDM) secrete measurable quantities of IFNα and IFNβ proteins in response to Mtb infection? While it is widely proven that macrophages activate type I IFN transcriptomic signalling, surprisingly, there is no direct experimental validation supporting the secretion of these proteins during Mtb infection, leaving this critical aspect of macrophage infection biology unresolved. Existing studies have demonstrated that mouse bone marrow-derived macrophages (BMDMs) and human monocytic cell lines secrete type I IFNs in response to Mtb infection[4-6], however, this has not been shown in primary human MDMs. To date, the available data have been limited to transcriptional evidence, with reports of IFN genes being upregulated during infection. Yet, the detection and quantification of the corresponding proteins remain unaddressed.

The widely studied H37Rv strain is fully virulent, whereas its attenuated counterpart, H37Ra, lacks key virulence factors (e.g, *phoP, mce1*, and *esx-1* secretion system components) that modulate immune evasion and intracellular survival [7, 8]. While both strains activate transcriptional type I interferon (IFN) signalling, their differential virulence may shape the magnitude and kinetics of cytokine secretion. H37Rv’s intact *esx-1* system was found to drive robust cytosolic DNA sensing via cGAS-STING, potentially amplifying IFNβ responses[9], whereas H37Ra’s lack of virulence may alter this balance.

This study therefore aims to fill a crucial gap in the literature. Using a commercial IFNβ ELISA kit and an ultrasensitive single-molecule array (Simoa) assay to detect IFNα, we provide for the first time, evidence that primary human MDMs secrete both IFNα and IFNβ proteins during a five-day course of Mtb infection and observed trends suggesting strain-specific modulation of IFN secretion.

## Methods

Peripheral blood mononuclear cells (PBMCs) were isolated from the buffy coats of healthy donors from the Irish Blood Transfusion Services using density-gradient centrifugation over Lymphoprep (StemCell Technologies). PBMCs were washed, resuspended at 2.5x10^6^ PBMC/ml in RPMI (Gibco) supplemented with 10% AB human serum (Sigma-Aldrich) and plated in non-treated tissue culture plates (Costar). Cells were maintained in humidified incubators for 7 days at 37°C and 5% CO2. Non-adherent cells were removed by washing every 2 days. Mtb H37Ra and H37Rv were obtained from The American Type Culture Collection (ATCC 25177TM; Manassas, VA) and propagated in Middlebrook 7H9 medium supplemented with ADC (Beckton Dickinson) to log phase. The multiplicity of infection (MOI) and donor variation in phagocytosis of Mtb was adjusted for by Auramine O staining, as described previously[10]. MDMs were infected at an MOI of 1-5 bacteria per cell and supernatants assessed 48 h and 120 h post infection. Supernatants were removed and IFNα quantified using the Simoa Pan-IFNα assay developed with Quanterix Homebrew kits as previously described [11]. IFNβ was quantified using an R&D DuoSet ELISA kit (Bio-techne) according to the manufacturer’s instructions. Statistical analysis was performed using a paired t-test to determine differences between groups with *p*<0.05 as significant.

## Results

Supernatants from H37Rv-infected macrophages exhibited a significant increase in IFNα levels at 48 h post-infection (mean 224.6 fg/ml) compared to uninfected controls. Similarly, H37Ra infection also led to a significant increase in IFNα levels at 48 h (mean 2220 fg/ml) relative to uninfected controls (Fig. 1A and B). IFNα levels persisted 120 h post-infection in both infection models however there was no significant difference between IFNα levels at both time-points. H37Rv infection significantly increased the secretion of IFNβ at both 48 h (mean 2.79 pg/ml) and 120 h (3.74 pg/ml) post infection vs. control, however H37Ra infection led to a significant increase in IFNβ levels only after 120 h of infection (mean 7.69 pg/ml).

**Figure 1.**
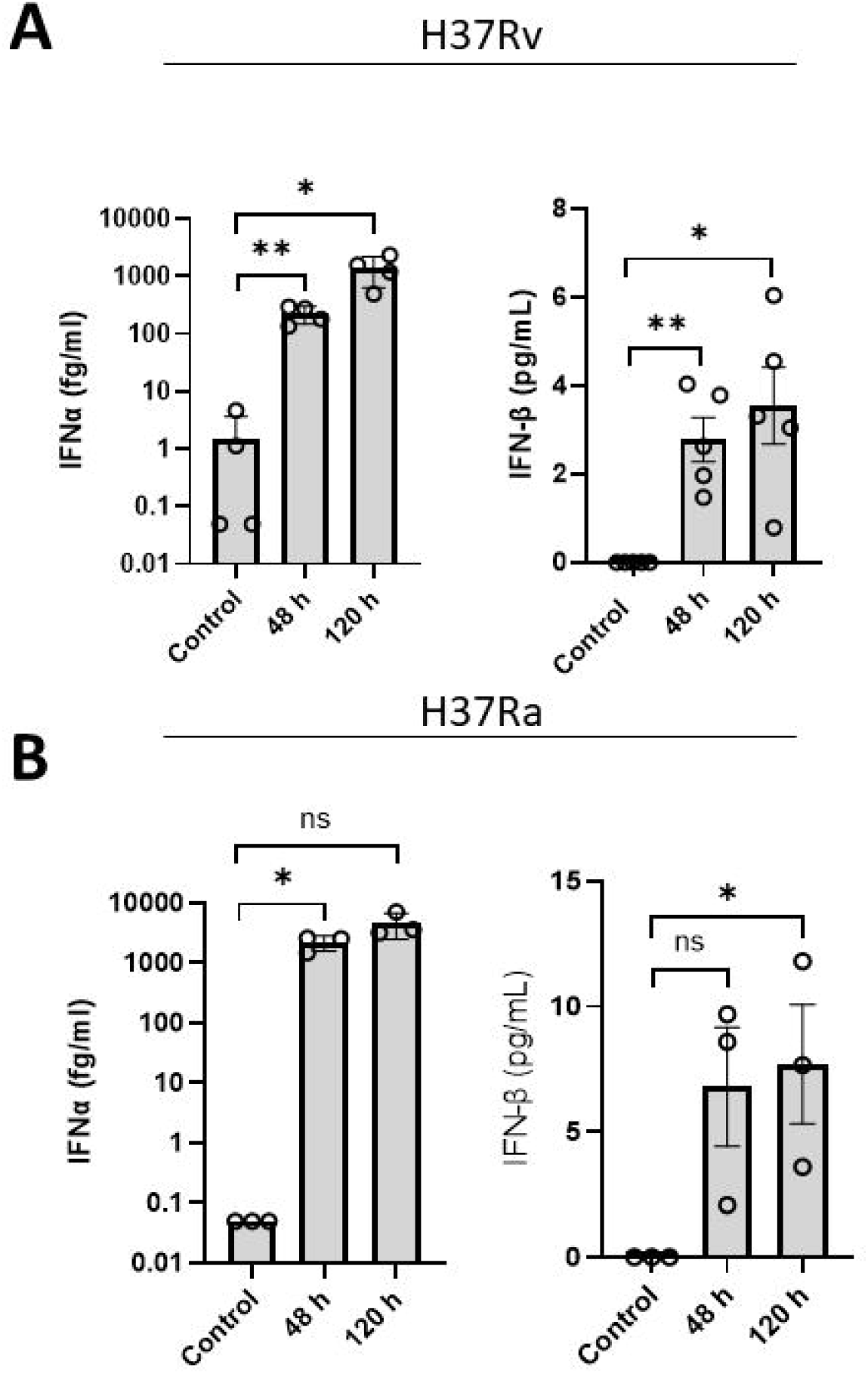
Primary human MDM secrete both IFNα and IFNβ during infection with H37Ra and H37Rv over a 120 h infection period. Human MDMs were infected with H37Rv (A) or H37Ra (B) for 48 h and 120 h at an MOI of 1-5 and supernatants removed to assess IFNα and IFNβ concentrations using the Simoa assay and ELISA, respectively. Data represent (A) n=4-5, (B) n=3.^*^p<0.05 and ^**^p<0.01 determined using a paired t-test.

## Discussion

The interaction between *Mycobacterium tuberculosis* (Mtb) and macrophages has been a focal point of TB research, with in vitro studies serving as an essential tool to dissect host-pathogen interactions. As mentioned previously, transcriptional signals for both IFNα and IFNβ exist, however their corresponding proteins have not been detected in primary human MDMs. In this study, we demonstrate, for the first time, that human MDMs secrete measurable levels of both IFNα and IFNβ proteins following a 5-day time course of Mtb infection, with secretion patterns influenced by the infection duration.

We utilised a simple, reproducible method of MDM differentiation based on plastic adherence of PBMCs in the presence of 10% human serum for seven days. This approach is widely used for its straightforward nature and minimal manipulation that yields MDMs of >90% purity without additional polarization stimuli. However, this method represents just one of many differentiation protocols in the field. A previous study compared two common differentiation methods—one involving M-CSF (macrophage-colony stimulating factor) and the other GM-CSF (granulocyte-macrophage colony stimulating factor)—highlighting how these variations can produce macrophages with distinct functional profiles[1]. The authors did in fact detect interferon stimulated genes 24 h post infection, which is line with other findings mentioned previously, but could not detect the protein using commercial methods. Thus, we don’t believe that the monocyte differentiation method is key in detecting type I IFNs, we do believe that the time point is crucial.

The majority of studies focus on 24 h time points to assess type I IFN release but have been unable to detect type I IFN protein and/or rely on transcriptional methods to confirm their presence. We observed that IFNα protein was detectable at 48 h post-infection and maintained these levels at 120 h. It could be argued that the presented concentrations of IFNα and β are functionally insignificant due to their strikingly low levels, however we would argue against this based on evidence that the induction of ISGs occurs as early as 6 h post-infection [12, 13] and strongly suggests that type I IFNs are being endogenously produced (although undetectable with current methods) and acting in both an autocrine and paracrine manner to activate ISG transcription. Furthermore, we would argue that these concentrations are functionally significant since their blockage using anti-IFN antibodies affect M.tb intracellular replication[5, 14]. We have also previously observed a significant positive correlation between equivalent plasma IFN protein concentrations and ISGs, strongly suggesting that fg/mL levels are indeed physiologically relevant [11]. Our findings emphasize that detectable type I IFN responses are time-dependent and suggest that shorter time points (<48 h) may fail to capture the full dynamics of these cytokines.

Our study also raises an important biological question: are these cytokines produced predominantly by Mtb-infected macrophages, or do bystander macrophages contribute to the observed secretion? Given the inherent heterogeneity of infection within macrophage cultures, single-cell analyses or spatial profiling techniques would be valuable for determining the cellular sources of type I IFNs during Mtb infection.

## Experimental Limitations

While this study provides valuable insights into type I IFN secretion, several limitations should be noted. We observed that H37Ra-infected macrophages exhibited higher mean IFNβ levels at 120 h compared to H37Rv-infected macrophages (7.69 vs. 3.74 pg/ml). While this comparison is limited by unmatched donors preventing statistical interpretation, the trend aligns with prior reports suggesting that attenuated Mtb strains may fail to fully suppress host defences due to mutations in immune evasion loci such as *phoP* [7, 15]. Conversely, H37Rv’s intact virulence factors could dampen late-phase IFN responses[15].

